# Identification of mutations associated with Macozinone-resistant in *Mycobacterium Tuberculosis*

**DOI:** 10.1101/2021.02.24.432818

**Authors:** Xi Chen, Yuanyuan Li, Bin Wang, Yu Lu

## Abstract

Macozinone is identified as a drug candidate and is currently under clinical development for the treatment of tuberculosis, but the mutations conferring resistance to Macozinone remain inadequately characterized. Herein, we investigated the Macozinone -resistance -associated mutations through selecting resistant isolates *in vitro*. Macozinone-resistant isolates were obtained through induction *in vitro*. The level of Macozinone -resistant strains was confirmed by MABA test. PCR sequencing analysis was carried out on dprE1 gene. Whole Genome Sequencing was performed to identify mutations associated with Macozinone -resistance. The totals of isolates obtained at Macozinone concentrations of 6.4ng/ml, 25.6ng/ml, 50ng/ml and 100ng/ml were 49, 20, 20 and 4 respectively. Among the 49 strains obtained by 6.4ng/ml Macozinone only one strain had C387S mutation in dprE1. C387S is only occurred in high-level resistant isolates (MIC > 500ng/ml). Meanwhile high-level resistance to Macozinone can occur in strains induced at 6.4ng/ml and the frequency of occurrence is low (1/49, 2.04%). The MIC_90_ of other strains except the strains carrying C387S mutation is at the same level (11.5ng/ml > MIC_90_ > 2ng/ml). The G61A or G248A mutations in dprE1 was discovered for the first time. Other gene mutations (rv0678, rrs, mbtF, rv2956, *et al*) were found in low-level resistant strains.

**Conclusions:** High-level resistant isolates can be produced at low concentration of Macozinone. C387S mutation in dprE1 is directly related to high-level resistance. There may be new mechanisms involved in Macozinone-resistance independent of dprE1 mutations.

---

Tuberculosis (TB) caused by *Mycobacterium tuberculosis* (*M.tb*) is an infectious disease that affects more than 10 million people worldwide every year, mainly in developing countries (1). It is estimated that approximately 1.7 billion people are infected with *M.tb* (1). Moreover, *M.tb* is still a global leading cause of death (1). The emergence and widespread of drug-resistant TB is now recognized as one of the most dangerous threats to global TB control (2). Patients with multidrug-resistant TB (MDR-TB), extensively drug-resistant TB (XDR-TB), and totally drug-resistant TB (TDR-TB) need long and expensive treatment (3–5). WHO approved anti-tuberculosis therapies can only treat less than 50% of MDR-TB and 30% of XDR-TB (6,7). There is an urgent need for the development of more efficient new drugs and shorter treatment regimens.

Macozinone (MCZ), previously known as PBTZ169, is currently undergoing Phase 1/2 clinical trails for the treatment of tuberculosis (8,9). MCZ had good bactericidal activity against multidrug-resistant tuberculosis strains and was effective against *M.tb* in replication phase (10–12). In the mouse infection model, MCZ showed excellent therapeutic effect. Other studies have also found that MCZ has synergistic effect with bedaquiline (10). MCZ targets the essential flavoenzyme DprE1 and blocks the synthesis of the cell wall precursordecaprenyl-phospho-arabinose (DPA) and provoking lysis of *M.tb* (10,13–19). Recent studies have confirmed that MCZ can form stable covalent complexes with DprE1 by targeting DprE1 enzyme (20). DprE1 plays a catalytic role together with DprE2 and catalyzes the epimerisation of decaprenyl-phospho-ribose (DPR) to DPA. DPA is the only donor of D-arabinose in mycobacteria (16–20). Arabinose polymers form DPA, which is one of the cell wall components of mycobacteria. The inhibition of MCZ on DprE1 eventually led to cell lysis and cell death (17).

It has also been verified that Cys387 in dprE1 is essential for the activity of DprE1 inhibitors (11). Cys387 plays an important role in covalent binding with MCZ and drug resistance occurs when the site is substituted by other amino acids (10,17,18). However, It is not clear whether the substitution of Cys387 residue in DprE1 is the single reason of MCZ-resistance. The characterization of MCZ-resistance is still unclear. In this study, the occurrence of resistance to MCZ similar to that of drug-resistant TB in clinical therapy was obtained through rising gradually MCZ concentration induction *in vitro*, so as to find out the mutations of drug resistance gene and to better understand the mechanisms of MCZ-resistance, which would be helpful for monitoring the resistance to MCZ and developing more accurate molecular tests in future clinical application.

## RESULTS

### Isolation of MCZ -induced isolates and Determination of the minimum inhibitory concentration (MIC)

The MIC_90_ of MCZ for *M.tb* H37Rv determined by MABA was 0.2ng/ml. To isolate induced MCZ-resistant strains, early stationary phase cultures of H37Rv were plated on 7H10 agar plates containing 0.1ng/mL MCZ. Isolated strains were obtained through several rounds of selection. After repeated MCZ susceptibility testing to rule out false resistance, 4 isolated strains obtained by MCZ concentration of 100ng/ml, 20 isolated strains obtained by MCZ concentration of 50ng/ml, 20 isolated strains obtained by MCZ concentration of 25.6ng/ml and 49 induced strains obtained by MCZ concentration of 6.4ng/ml were selected. Only 4 strains survived when the MCZ concentration in the 7H10 medium increased to 100ng/ml. The MIC_90_ values of above isolated strains were measured respectively (shown in table 1). Because there is no clinical MCZ-resistance break point now, MIC_90_ value of isolates higher than 2 ng/ml is defined as MCZ resistance (>10 × MIC_90_ of H37Rv), and MIC_90_ value higher than 500ng/ml is defined as MCZ high-level resistance in our study. Finally, a total of 4 high-level resistant strains were obtained, among which 3 strains (S1-1,S2-1,S3-1) isolated from the medium with MCZ concentration of 100ng/ml and 1 strain (S4-4) isolated from the medium with MCZ concentration of 6.4ng/ml. The emergence of S4-4 showed that high-level resistance to MCZ can occur in stains induced at low-level drug concentration. Meanwhile by the induction of 6.4ng/ml MCZ, the frequency of occurrence of high-level resistance (1/49) is low which showed that it is difficult to produce high-level resistance strains to MCZ. Although strains S4-1, S4-2 and S4-3 were also obtained by culture step by step, they were thought to be the same strain because of their same genome sequence by sequencing analysis of the whole genome.

**Table 1.**
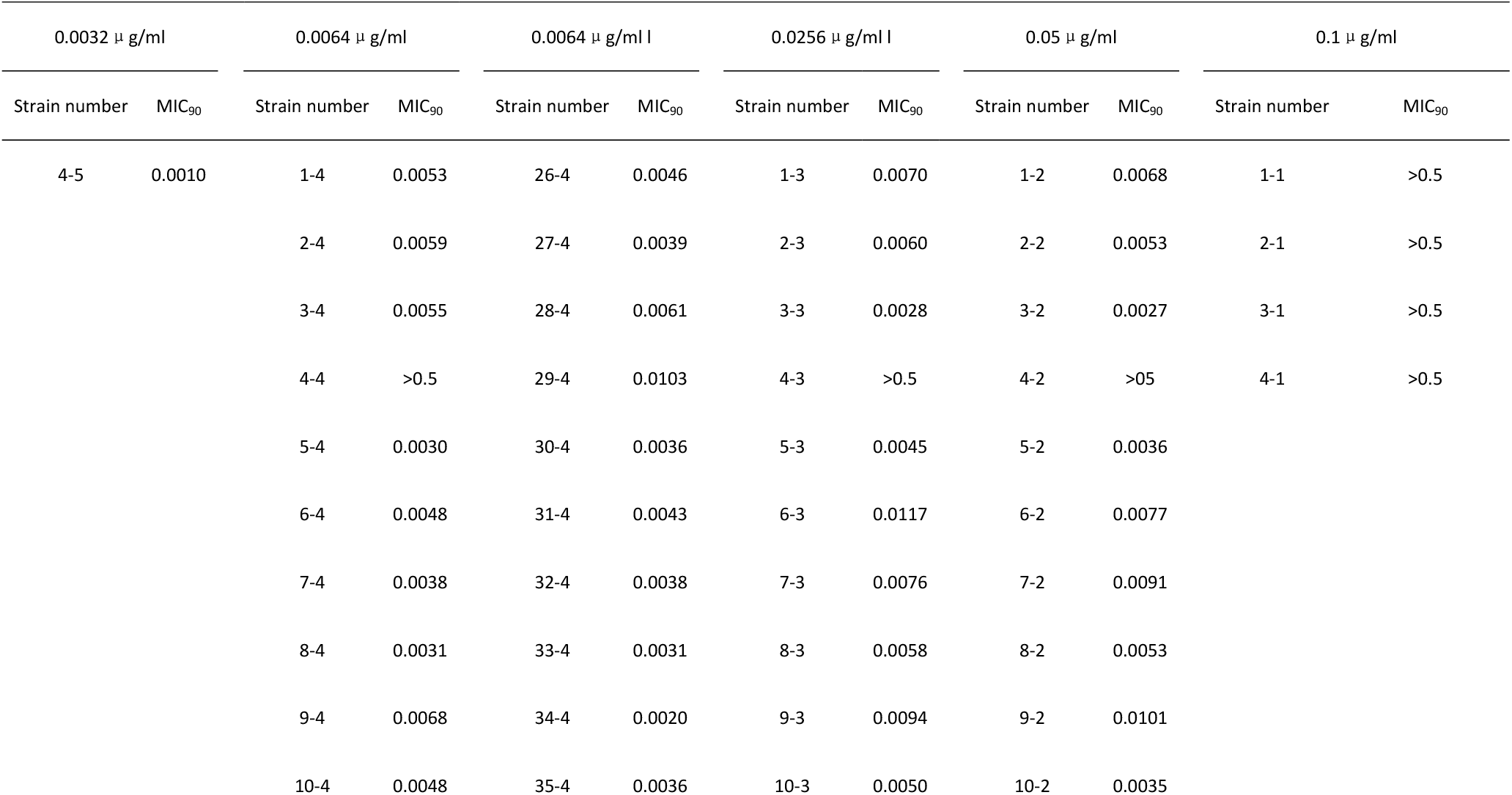
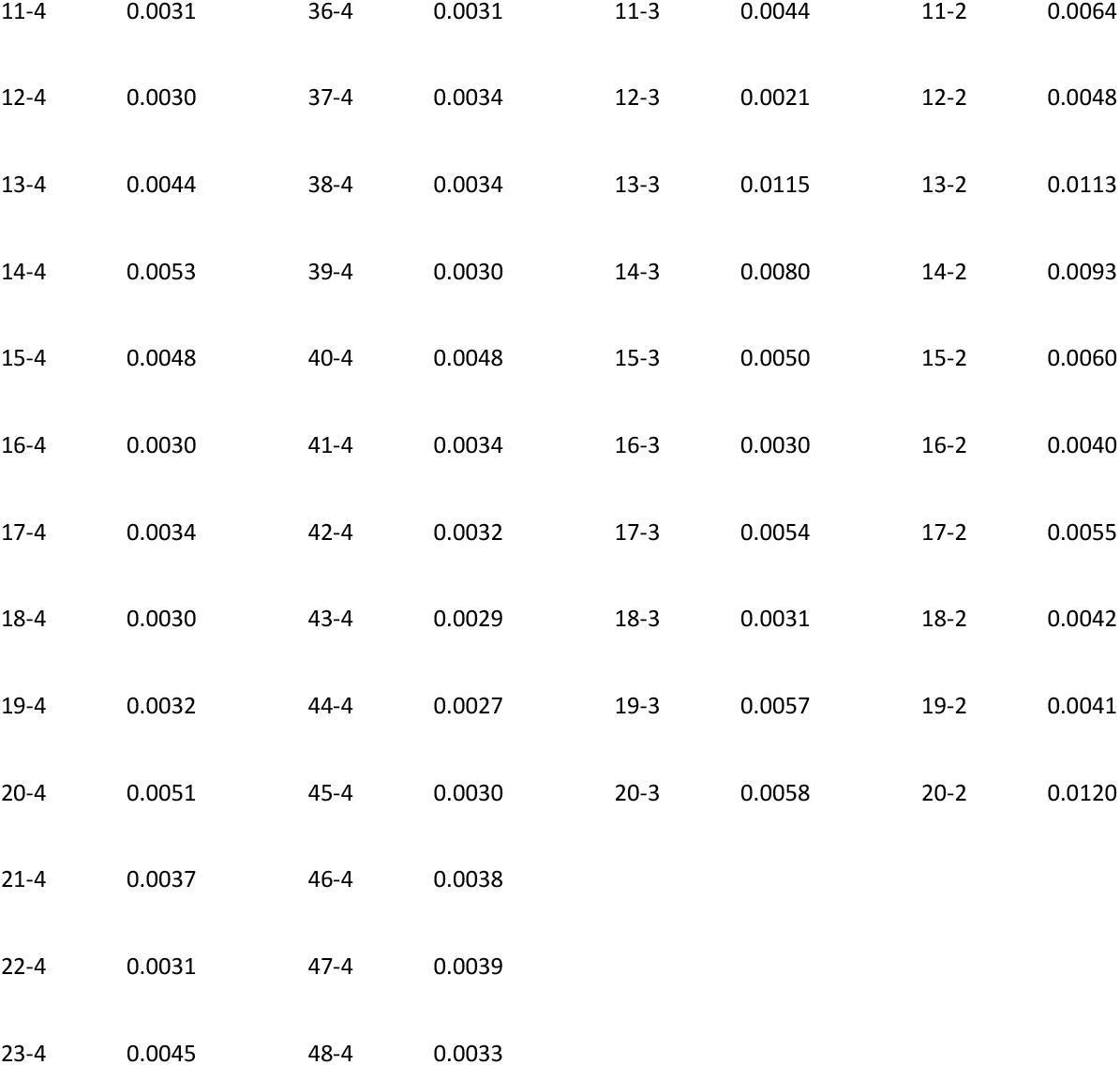
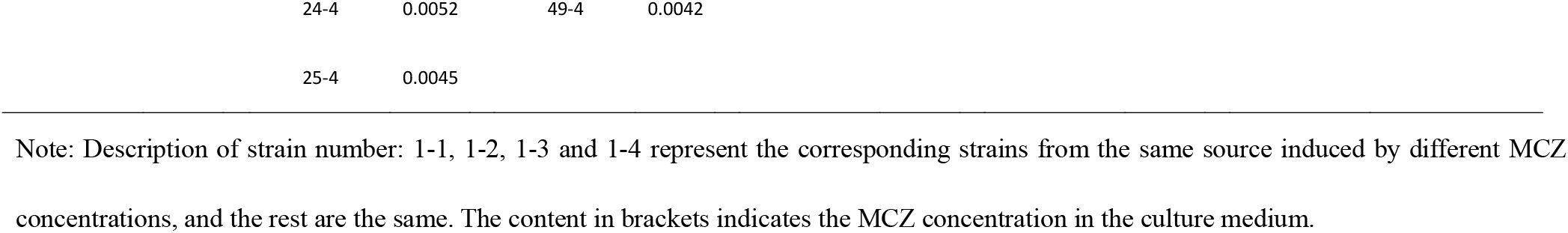
MCZ sensitivity detection of induced strains (MIC_90_, μ g/ml)

Except for 4 high-level resistant strains, the rest of isolated strains had lower and similar MIC_90_ value (11.5ng/ml > MIC90 >2ng/ml), which were more than 10-fold MIC of Wide-type H37Rv. For these low-level MCZ-resistant strains, even the concentration of MCZ in 7H10 solid medium rised to 50ng/ml, MIC_90_ were no longer increased (Table 1).

### dprE1 sequencing

We isolated genomic DNA from the 49 MCZ-induced strains at the concentration of 6.4ng/ml and performed PCR to amplify 674 bp DNA fragment of dprE1 gene. Only strain S4-4 had C387S mutation (tGc/tCc, 1/49, 2.04%) whose MIC was > 500ng/ml, whereas other 48 strains had no C387S mutation whose MIC was between 2ng/ml and 10.3ng.ml. This result suggested that there is an obvious correlation between C387S mutation in dprE1 and MCZ-resistance level. In order to know whether the mutation of C387S may occur under the lower MCZ concentration, Cys387 in S4-5 obtained by inducing at MCZ concentration of 3.2 ng/ml had also been detected. We found that S4-5 had no C387S mutation. Considering C387S mutation can occur in strain S4-4 at the concentration of 6.4ng/ml, this result means low concentration MCZ induction can produce high-level drug-resistance and C387S mutation in dprE1 only occurs in high-level MCZ-resistant strains.

### Sequencing Analysis of the Whole Genome of H37Rv MCZ-resistant Strain

To explore the development of MCZ resistance *in vitro* and to compare the genes changes before and after drug induction, a total of 9 MCZ-resistant strains were subjected to whole genome sequencing using Illumina HiSeq platform. The 9 strains were S2-1, S2-2, S2-4, S3-1, S3-2, S3-4, S4-1, S4-4 and S4-5. Among these isolates S2-1, S3-1, S4-1, S4-4 were high-level resistance strains (MIC > 500ng/ml), whereas others were low-level resistant strains (MIC > 2ng/ml). We found that S4-1 and S4-4 have C387S mutation (tGc/tCc) which is consistent with the previous PCR results. Another 2 high-level resistant strains (S2-1, S3-1) also had C387S mutation (Tgc/Agc, tGc/tCc). It is interesting to note that 3 strains (S2-1, S2-2 and S2-4) had G248A mutation in the dprE1 gene (gGc/gCc) and 3 strains (S3-1,S3-2 and S3-4) had G61A mutation in dprE1 (gGg/gCg).

Comparative genome sequence analyses of the 9 strains revealed that they all had missense mutations in 3 genes, fadD2 (R137P, 9/9, 100%), phoT (Q153R, 9/9, 100%) and rv1725c (E28G, 9/9, 100%); they also had frame shift mutation in SenX3 gene (animo acid 330 site, C/CG, 9/9, 100%) (Table 2). Isolates S2-1, S2-2 and S2-4 also had A99V substitution in rv0678 (3/9, 30%), Q1026H substitution in rrs (3/9, 30%) and A108T substitution in rv2956 (3/9, 30%). A270G substitution in mbtF appears in isolates S2-1, S2-2, S2-4, S4-1, S4-4 and S4-5 (6/9, 66.7%). Isolates S3-1 had a S290P mutation in rv3230c (1/9, 11.1%). Intergenic region of rv1706A-rv1706c mutation was only found in S3-1 and S3-2 (2/9, 22.2%). S4-1 and S4-4 had also identical aofH mutation G383S (2/9, 22.2%). Our findings suggested that these mutations independent of dprE1 may be related to the increased sensitivity of MCZ induced low-level resistant strains.

**Table 2.** Genome sequencing of some strains

| Strain number | genome sequencing |
| --- | --- |
| 2-1 | **dprE1(C387S)**, rv0678(A99V),rrs(Q1026H),mbtF(A270G),rv2956(A108T),**dprE1(G248A)**,fadD2(R137P),phoT ( Q153R ) ,rv1725c (E28G),SenX3 (frameshift) |
| 2-2 | rv0678(A99V),rrs(Q1026H),mbtF(A270G),rv2956(A108T),**dprE1(G248A)**,fadD2(R137P), phoT(Q153R),rv1725c (E28G) ,SenX3 (frameshift) |
| 2-4 | rv0678(A99V),rrs(Q1026H),mbtF(A270G),rv2956(A108T),**dprE1(G248A)**,fadD2(R137P), phoT(Q153R),rv1725c (E28G) ,SenX3 (frameshift) |
| 3-1 | **dprE1(C387S)**,**dprE1(G61A)**,fadD2(R137P),phoT (Q153R),rv1725c(E28G), ,SenX3 (frameshift), rv3230c(S290P),intergenic region |
| 3-2 | intergenicregion,**dprE1 (G61A)**,fadD2(R137P),phoT(Q153R),rv1725c(E28G) ,SenX3 (frameshift) |
| 3-4 | **dprE1 (G61A)**,fadD2 (R137P), phoT ( Q153R ) , rv1725c (E28G) ,SenX3 (frameshift) |
| 4-1 | **dprE1(C387S)**,mbtF(A270G),aofH(G383S),fadD2(R137P),phoT (Q153R), rv1725c (E28G) ,SenX3 (frameshift) |
| 4-4 | **dprE1(C387S)**,mbtF(A270G),aofH(G383S),fadD2(R137P), phoT (Q153R), rv1725c (E28G) ,SenX3 (frameshift) |
| 4-5 | mbtF(A270G),fadD2(R137P), phoT (Q153R), rv1725c (E28G) ,SenX3 (frameshift) |

Note: Description of strain number: 1-1, 1-2, 1-3 and 1-4 represent the corresponding strains from the same source induced by different drug concentrations, and the rest are the same.

## DISCUSSION

MCZ is a bactericidal benzothiazinone that inhibits the essential flavoprotein DprE1 by forming acovalent bond with the active-site Cys387 residue, thus preventing the synthesis of decaprenyl-phosphoryl-ribose. MCZ resistance in *M.tb* is caused by mutations in dprE1 gene. A previous study that employed genetic approaches confirmed that rv3790 gene in BTZs-resistant *M. tb* mediated increased resistance and mutants harbor missense mutations in this gene (14). DprE1(Rv3790) is a drug target of MCZ (8,14,15,17). It had been verified that Cys387 is essential for the activity of DprE1 inhibitors by method of promiscuous site-directed mutagenesis to introduce other codons at Cys387 into dprE1 of *M.tb* (18). The Cys387 residue of DprE1 is highly conserved in orthologous enzymes in mycobactria and substitution of Cys387 with alanine or serine in *M.avium*, *M.intracellulare* and *M.abscessus* may weaken the binding affinity between the drug and the target, thus showing the natural drug resistance of these NTM strains to MCZ (13,18,19).

In this study we obtained *M.tb* MCZ-resistant isolates *in vitro* through H37Rv growing on agar plates containing MCZ from low concentration to high concentration. We found that C387S mutation in dprE1 only occurred in high-level MCZ resistant isolates which is consistent with previous report in which Cys387 position harboring alleles showed high MCZ resistance (18). Drug resistance occurs spontaneously in *M.tb* at a different rate for each drug. For example, mutations resulting in resistance to RIF occur at a rate of 10^−10^/cell division, compared with 10^−7^–10^−9^ for INH (21). The frequency of Cys387 amino acid position in dprE1 harboring alleles is middle (<10^−8^) (8). This kind of middle spontaneous mutation rate (<10^−8^) may explain the emergence of high-level resistant strains to MCZ by the induction at low drug concentration of 6.4ng/ml in our research. This phenomenon also means that it may be one-step mutation of dprE1 gene in strain S4-4, rather than the accumulation of mutations causing high-level resistance. The result also suggested that selective pressure of drug on strain is obvious and attention should be paid to the possibility of drug resistance of *M.tb* at low MCZ blood drug concentration caused by irregular or low-dose administration in future clinical application. Especially for the patients with lung cavities the risk of drug resistance may be higher because lung cavities frequently contain 10^7^ resistant bacilli emerging naturally without antimicrobial pressure, then the use of anti-mycobacterial drugs is prone to select the resistant population (22–24).

In addition, G61A and G248Amutations in dprE1were discovered for the first time. They appeared not only in high-level resistant isolates but also in low-level resistant isolates. So their contribution in the MCZ-resistance remained to be determined in the following research. We obtained a number of low-level resistant isolates whose decreased sensitivity to MCZ isn’t illustrated by dprE1 mutation. Considering the mutations in known gene can’t explain MCZ resistance in these isolates the possibilities must be considered that there may be other gene mutations conducive to MCZ resistance. In our research we indeed found some mutations in several genes, such as Rv0678, Rv3230c *et al*. But these genes mutation were detected in only 9 strains, it is difficult to speculate that they are related to low-level resistance to MCZ. More gene mutations that may be related to MCZ resistance would be found by expanding the sequencing number of induced isolates in future research.

It is confirmed that resistant bacteria is prone to fitness cost, taking the form of the decreased viability and virulence compared to the wild-type strain in the absence of antibiotics. Fitness cost is common in bacteria (25–30). Drug-resistance of bacteria often changes some structures such as DNA helicases, ribosome, RNA polymerase, cell wall, and so on. This kind of change often leads to the decrease of bacterial growth speed and virulence, which means the acquisition of drug-resistance needs to pay an adaptive cost (31,32). Therefore, the occurrence of drug-resistant mutations is a double-edged sword. Similar phenomena also exist in INH-resistant or RIF-resistant *M.tb* (33,34). However, drug-resistant bacteria can restore their fitness through evolution, a process called “compensatory evolution”. One of the reason of compensatory evolution is that drug-resistant bacteria can further accumulate other specific mutation through evolution change. Drug-resistant *M.tb* also can accumulate compensatory mutations and may recovery the fitness, which is helpful for the spread of drug-resistant *M.tb* in the population (35–40). In our study, all of the 9 strains had nonsynonymous mutations in four non-drug-resistant related genes (fadD2, phoT, rv1725c and SenX3) changed significantly during drug induction, suggesting a possible compensatory evolutionary mechanism. On the other hand, the relationship between bacterial drug resistance and its fitness provides a new insights into analyzing the causes of drug resistance and exploring how to reduce the possibility of drug resistance.

There are limitations that must be taken into account. Firstly, there is no clinical strain yet, and the induced strains were taken as the research object; Secondly, less drug-resistant strains were obtained. The number of whole genome sequencing strains was small. However, this study still had clinical significance: (1) Insufficient drug application may lead to high-level resistance of MCZ in future clinical treatment; (2) C387S mutation in dprE1 can be used as a marker for identifying high-level resistant strains.

In conclusion, we confirmed that C387S mutation in dprE1 is directly related to the emergence of high-level MCZ-resistant strains which can be used as a molecular marker to detect MCZ resistance in future clinic application and identified that there are other drug resistance mechanisms, which are related to low-level resistance. The mechanism of low-level resistance can be clarified by genome sequencing of more kinds of induced strains in future work.

## M ATERIAS AND METHODS

### Bacterial strains and growth conditions

*M.tb* H37Rv (ATCC 27294) is frozen in the department of pharmacology of Beijing tuberculosis chest tumor research institute. Subcultures in 7H9 broth were grown up to the logarithmic phase and bacterial suspension was filtered by 8 μm filter membrane to obtain bacterium suspension. Single bacterial suspension was inoculated into 7H10 culture plate after being diluted ten-fold, colony forming unit (CFU) of the single bacterium suspension is determined after 4 weeks. Viable bacterial were preserved in 1.0 ml aliquots frozen at −80°C. For each experiment, one aliquot was diluted in 7H9 broth to obtain a desirable concentration of bacteria.

### Susceptibility testing

MCZ susceptibility testing was carried out using MABA method to determine minimum inhibitory concentration (MIC_90_) as previously described (41). Briefly, prepare *M.tb* single colony suspension: *M.tb* is inoculated into 7H9 broth containing 10% OADC and 0.05% tween-80, cultured at 37 °C and 5% CO2 for 2~3 weeks to logarithmic growth phase. The isolate in the log phase was diluted to 1×10^6^cfu/ml in 7H9 broth containing 10% OADC. MCZ was serially diluted twofold in 100 μl 7H9 broth, and the final concentrations were 0.5, 0.25, 0.125, 0.0625, 0.03125, 0.016, 0.008, 0.004, 0.002, 0.001, 0.0005, 0.00025 and 0.000125μg/ml. Then, 100 μl bacterial suspension was inoculated to each well of microplates. The final volume of each well was 200 μl, containing 100 μl of MCZ solution and 100 μl of bacterial suspension. Furthermore, MCZ -free wells were used as positive control to determine the time for adding alamarBlue. Microplates were incubated at 37 °C for 7 days. 20 μl alamarBlue mixed with 12.5 μl 10% Tween-80 was added to the Microplates. The minimum inhibitory concentration (MIC_90_) value was defined as the lowest MCZ concentration that inhibited bacterial growth and prevented a color change.

### Protocol for MCZ induction

The method of *in vitro* induced resistant to MCZ is as shown in figure 1. Briefly, MCZ was dissolved in DMSO at a stock concentration of 1mg/mL and incorporated into 7H10 agar plates containing OADC at concentrations of 0.1ng/mL. 5×10^5^ cfu/ml H37Rv were inoculated on the above 7H10 agar plate. Isolates that grew on the MCZ containing plates after 3~4 weeks incubation at 37°C were picked and inoculated on the 7H10 agar plates containing OADC at concentrations of 0.2ng/mL. By repeating above steps isolates were obtained from 7H10 agar plates containing MCZ from 0.1ng/mL - 100ng/mL. All the single colonies were collected and stored in 7H9 broth at −80°C respectively. Then isolates were grown in 7H9 broth for confirming MCZ resistance phenotype by MABA method. Wild type H37Rv was included as a drug susceptible control strain for the MCZ susceptibility testing. In the MCZ susceptibility testing isoniazid (INH) and rifampicin (RIF) was used as control drug.

**Figure 1.**
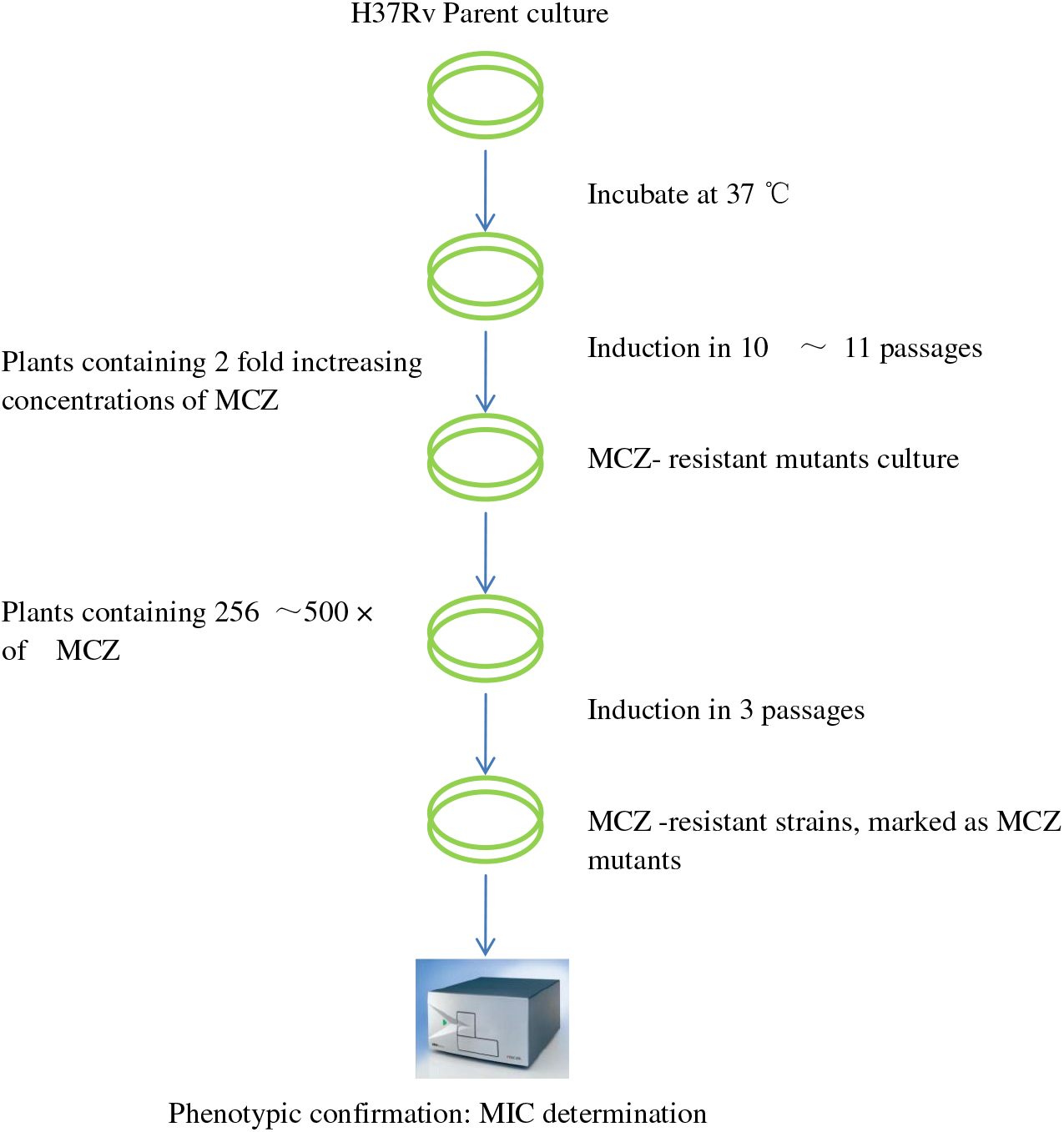
The method of *in vitro* induced resistant to MCZ.1

### Extraction of genomic DNA of MCZ -induced strains

Single colonies obtained from 7H10 agar plates with MCZ concentration of 3.2 ng/ml ~ 100ng/ml were selected. Genomic DNA was extracted using the Wizard ® Genomic DNA Purification Kit (Promega, American) according to the manufacturer’s instructions.

### PCR amplification of dprE1 and Detection of mutations associated with MCZ resistance

Strains obtained from the 7H10 agar with MCZ concentration of 6.4ng/ml were selected to carry out PCR sequencing analysis on dprE1 gene. The dprE1 PCR was performed using forword primer (5’-CTCACGCAGTTCTACCATCCG-3’) and reverse primer (5’-TCCAGGGCGTCAAAGTCG-3’) as described. Briefly, genomic DNA from MCZ-induced strains was used as templates for PCR as follows: heat denaturation at 94 °C 5 min followed by 30 cycles of 94 °C 1 min, 55 °C 1 min, 72°C 1 min followed by extension at 72 °C for 10 min. The PCR reaction was then cooled to 4°C. The dprE1 PCR products were then sequenced by ABI 377 DNA sequencer at Beijing ReboXingke Biotechnology Co., Ltd., and the dprE1 sequences from different isolates were compared against the wild type dprE1 sequence of H37Rv to identify potential mutations in the dprE1 gene.

### Whole genome sequencing

The genomic DNA for whole genome sequencing was isolated as previously described. Sequencing libraries were prepared using the Nextera XT Sample Prep Kit (Illumina, San Diego, CA, USA) according to the manufacturer’s protocol and sequenced on Illumina Hiseq and BGISEQ-500 platforms. The sequencing depths is 50-fold coverage. High-quality reads were aligned to the *M.tb* H37Rv (GenBank NC000962.3) reference sequence using Bowtie2. Single nucleotide polymorphisms (SNPs) and InDels (insertion and deletion) were detected using SAMtools. In order to eliminate the genomic differences of MCZ-resistant islolates and H37Rv in our analysis, SNPs and InDels were further annotated for gene locus and mutation types with the nearest coding sequences.

Synonymous mutations and PE/PPE mutations within coding sequence were removed in the final analysis to focus on mutations that are most likely involved in MCZ-resistance. The whole genome sequencing analysis was carried out in Shanghai Bo-hao Biotechnology Company.

## ACKNOWLEDGEMENTS

This work was supported by National Science and Technology Major Project of China (2019ZX09721-001-007-003).

The funders had no role in study design, data collection and interpretation, or the decision to submit the work for publication.

We have no conflict of interest to declare.

